# Defining a Midgestational Window for *In Utero* Genome Editing of the Fetal Murine Cortex

**DOI:** 10.64898/2026.04.28.721509

**Authors:** Cameron R. Jackson, Máté Borsos, Nathan Appling, Carrie R. Jackson, Gerard M. Coughlin, Viviana Gradinaru

**Affiliations:** Division of Biology and Biological Engineering, Caltech, Pasadena, CA 91125; Howard Hughes Medical Institute, Caltech, Pasadena, CA 91125

## Abstract

Congenital disorders of cortical development arise from genetic lesions that disrupt neurogenesis and neuronal migration. Unfortunately, tools to model or correct these defects before birth are limited. Here we establish a platform for systemic *in utero* gene delivery and genome editing in the mouse cortex at midgestation. By microdissecting a uterine window over the vitelline vein at embryonic day 12.5 (E12.5), we achieve fetal circulation access, enabling robust AAV-mediated transduction of the central nervous system (CNS) while reducing off-target expression in peripheral organs. Barcoded capsid screens reveal that AAV9 exhibits developmental stage-dependent tropism, with higher CNS penetrance and lower liver transduction at E12.5 than at E15.5. Leveraging this window, we provide a proof-of-concept of efficient cortical editing, using Cre-lox and CRISPR/Cas9 strategies to recapitulate prenatal reeler-like cortical misordering phenotypes following *Reln* knockout. We further use homology-directed repair to demonstrate precise genome modification, epitope-tagging the endogenous *Reln* and *Actb* loci, and installing a human-derived pathogenic allele of *PDHA1*. Importantly, we show that edited cells span neural progenitors and differentiated neurons across the cortex and hippocampus. These results define a permissive midgestational window for prenatal genome editing, providing a platform for functional modeling of congenital CNS disorders and exploration of early therapeutic interventions with minimized peripheral exposure.

## Introduction

Congenital disorders comprise a diverse group of conditions that arise during fetal development or at birth, leading to structural or functional abnormalities with lifelong consequences [1]. Among these, disorders affecting the central nervous system (CNS) are particularly devastating, as the brain undergoes rapid and highly coordinated development beginning shortly after fertilization and continuing into adolescence and adulthood [2, 3, 4]. The brain grows more rapidly *in utero* than at any other stage of life, rendering early neurodevelopment uniquely susceptible to genetic and environmental perturbations [5,6]. CNS abnormalities occur in nearly 1 in 100 live births, with malformations of the cerebral cortex representing a major subset of these conditions [7, 8].

Malformations of cortical development (MCDs) frequently arise from genetic variants affecting key neurodevelopmental genes [9]. A 2024 retrospective study of 32 prenatally diagnosed MCD cases identified single nucleotide or copy number variants in 22 affected fetuses, revealing the predominance of discrete genetic lesions in these disorders [10]. Notably, 31 of the 32 pregnancies were terminated, highlighting the absence of effective prenatal treatment options and the limited ability to predict postnatal outcomes. These findings underscore the urgent need for strategies that enable both functional modeling and therapeutic intervention during fetal brain development.

Prenatal gene therapy offers a compelling opportunity to intervene before irreversible neurodevelopmental defects are established [11]. Treating the fetus at early developmental stages has shown therapeutic potential across multiple mammalian systems [12]. Systemic delivery approaches have successfully accessed diverse fetal organs and cell types in both mouse and human, including in recent clinical studies in which enzyme replacement therapies delivered via the umbilical vein enabled prenatal treatment of an otherwise lethal lysosomal storage disorder [13]. These examples illustrate the feasibility and potential impact of treating disease at its developmental origin rather than after birth.

Recombinant adeno-associated viruses (AAVs) have emerged as leading vehicles for *in utero* gene therapy due to their favorable safety profile, low immunogenicity, and customizable genetic cargo [14, 15]. AAVs can deliver transgenes, regulatory elements, and gene editing machinery with minimal unintended genomic integration [16, 17]. Importantly, single-dose AAV administration is well tolerated by the fetal immune system, and the small size of the fetus enables efficient systemic delivery using substantially lower vector doses than those required in postnatal or adult settings [18]. Despite these advantages, significant challenges remain. Efficient and selective delivery to the developing CNS remains difficult, and systemic administration of AAVs often results in substantial off-target liver transduction, limiting therapeutic efficacy and raising safety concerns [14, 19].

To date, most AAV-based prenatal therapies have relied on gene replacement strategies, in which therapeutic efficacy depends directly on transduction efficiency in target cells [20]. In the developing fetus, however, rapid cell division leads to dilution of episomal viral genomes, leading to diminished transgene expression [21, 22]. In contrast, genome editing offers the potential for permanent genetic modification, particularly when targeting proliferative progenitor populations early in development [23, 24, 25]. CRISPR/Cas9-based *in utero* genome editing has been demonstrated in the fetal mouse, but reported editing efficiencies in the brain remain low when editors are delivered systemically using AAV9 at or after embryonic day 14.5 (E14.5) [26]. As in juvenile and adult animals, the liver captures a substantial fraction of systemically delivered virus at these stages, limiting CNS access [27]. These observations suggest that developmental timing, rather than vector choice alone, may be a critical determinant of successful prenatal genome editing in the brain.

Midgestational stages, between E11 and E15 in the mouse, encompass critical transitions in both hepatic and cortical development. During this window, the fetal liver gains functionality, and neural progenitor populations undergo rapid expansion and migration [28, 29]. Early midgestation (E10-E12.5) marks the decline of dominating stemness-associated gene expression in the cortical neuroepithelium, followed by increasing neuronal differentiation, deep layer cortical lamination, and fate specification [30, 31, 32]. Access to the embryo during this period is therefore particularly attractive for genetic intervention targeting cortical progenitors. However, systemic AAV delivery at these stages is technically constrained. Placenta-crossing vectors capable of achieving reliable fetal CNS delivery have not yet been developed, rendering maternal injection ineffective [33]. Direct injection into the embryonic circulation circumvents this limitation but has historically been restricted to later gestational stages, as uterine opacity and tissue density prior to E14 prevent visualization of the murine vitelline vein using standard techniques [34].

Advancing safe and effective *in utero* genome editing in the developing cortex has broad implications for both therapy and disease modeling. From a translational perspective, editing the fetal genome of wild-type mice would enable rapid functional evaluation of pathogenic variants identified during prenatal screening, providing critical insight into disease severity and potential postnatal outcomes. Further, efficient genome editing in the embryonic cortex would enable rapid modelling of rare human neurodevelopmental disorders in mice within weeks. Coupling refined surgical access with modern genome editing tools thus offers a powerful framework for investigating cortical development, modeling congenital disorders, and exploring early therapeutic intervention strategies. Toward this goal, we integrate uterine microdissection-based vascular access with systemic AAV delivery to define a permissive early midgestational window for prenatal gene delivery and genome editing in the mouse cortex.

## Results

### Limited capsid-dependent CNS tropism at E15.5 suggests gestational permeability constraints

To identify the optimal AAV capsid for embryonic CNS delivery, we injected a barcoded pool of seven capsids, including AAV9, AAV-DJ, AAV1, AAV6, CAP-B10, PHP.eB, and X1.1, in equal concentrations into E15.5 embryos [37, 38, 39]. Normalized vector abundances were quantified across CNS (cortex and caudal brain) and non-CNS tissues, including liver, at both DNA and RNA levels. Brain-biased variants, engineered from the AAV9 backbone in adult mice, PHP.eB, CAP-B10, and X1.1 showed no statistically significant differences from AAV9 in either CNS or liver transduction at the DNA or RNA level (all p > 0.05) (**Figure S1**).

### Uterine microdissection enables E12.5 systemic access to the mouse embryo

To systemically deliver AAVs *in utero* at midgestational stages, we identified the vitelline vein as a microinjection target, based on previous *in utero* studies [34]. The vitelline vein conveys yolk sac blood to the hepatic primordium, where it helps to establish the hepatic sinusoids and portal circulation. This blood ultimately enters the fetal systemic circulation via intrahepatic pathways and the ductus venosus [40, 41]. The vitelline vein is visibly accessible for transuterine injection from E14 on. However, prior to E14 the uterus is opaque and dense around the smaller, midgestation embryos, thus complicating access for systemic injection. To access the embryo at an earlier stage, we dissected a window into the uterus directly over the unpaired vitelline vein (**Figure 1A, 1B**). This technical advancement enabled *in utero* systemic delivery to the E12.5 embryo (Supplementary video 1). Fluorophore-conjugated dextran dye injections at E12.5 revealed robust delivery to the developing brain, heart, liver, lungs, and gut. Injection at E15.5 resulted in high uptake of dye into the liver, relative to the brain and other organs (**Figure S2**).

**Figure 1:**
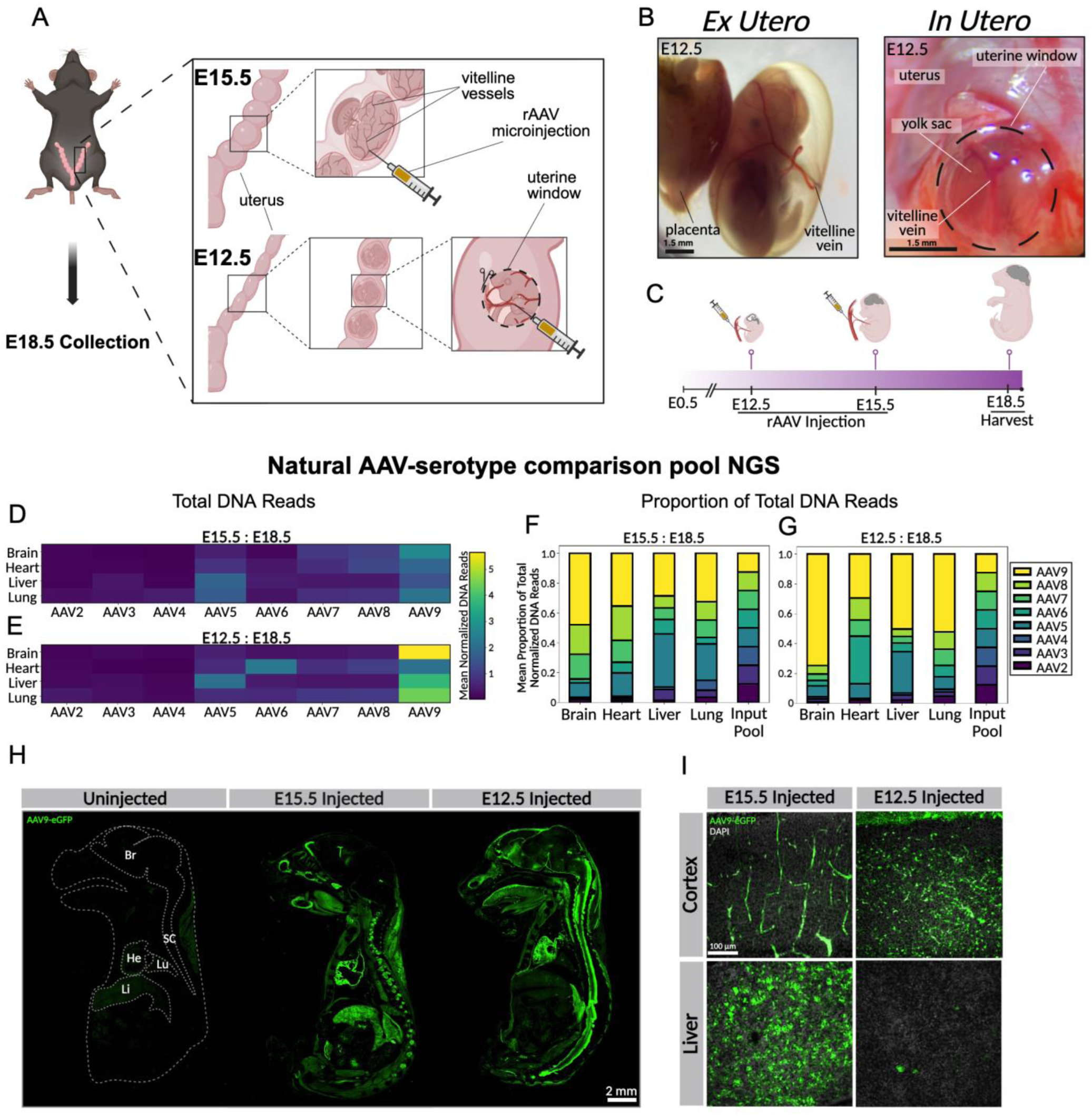
Systemic AAV9 delivery via uterine window enables robust CNS gene delivery at E12.5. **A)** Schematic of surgical techniques. E15.5 injection is performed through the uterus into the vitelline vein of the embryo. E12.5 injection requires microdissection of the opaque uterus to access the vitelline vein, followed by AAV microinjection. **B)** E12-E12.5 embryo within 1 hour of extraction and E12.5 embryo inside the uterus with yolk sac, uterus, and vitelline vein visible. **C)** Schematic illustrating timeline of *in utero* AAV administration at E12.5 and E15.5, with embryo harvest at E18.5. **D-E)** Barcoded AAV library NGS analysis showing age-dependent enrichment of AAV9 in the brain (n = 3 embryos per stage). Vector genome counts were normalized to the relative abundance of each capsid in the input barcoded AAV library. **F-G)** Stacked bar plots depicting the mean proportion of normalized reads per serotype at each developmental stage, calculated from the normalized vector genome counts shown in D-E, demonstrating a shift in recovered capsid composition with increased AAV9 representation in the brain after earlier stage injection. **H)** Sagittal sections of whole E18.5 embryos showing prenatal distribution of transgene expression following AAV9-mediated gene delivery at E12.5 or E15.5, visualized by anti-eGFP immunostaining. An uninjected control embryo is included. Anatomical landmarks are labeled as Br (brain), SC (spinal cord), He (heart), Lu (lung), and Li (liver). **I)** Representative higher-magnification images of cortex and liver from E18.5 embryos following AAV9 injection at E12.5 or E15.5, highlighting a developmental-stage-dependent shift in tissue tropism characterized by increased cortical transgene expression and reduced liver expression after E12.5 delivery (n = 3 embryos per stage).

### AAV9 shows developmental stage-dependent CNS tropism with greater penetrance at E12.5

To identify an ideal capsid for AAV-mediated *in utero* gene therapy, we assembled a barcoded pool of natural serotypes, selected based on structure and tropism diversity in mouse and other species. Anticipating differences in tropism across gestational stages, we delivered the natural serotype pool intravenously to embryos at stages E12.5 and E15.5 and harvested at E18.5, allowing 144- and 72-hour infection periods, respectively (**Figure 1C**). NGS analysis of recovered DNA from barcoded viral genomes revealed brain enrichment of AAV9, versus other serotypes. Furthermore, the brain enrichment of AAV9 was greater with E12.5 injection than with E15.5. These results suggest that AAV9 can provide efficient CNS access when injected at E12.5 (**Figure 1D, 1E**). Among all natural AAV serotypes included in the pool, AAV9 was the only capsid to display a consistent developmental difference in CNS tropism. In the E12.5-injected brain, AAV9 accounted for a mean of 73.25% of total normalized vector genomes, compared to 43.48% in the E15.5-injected brain (**Figure 1F, 1G**), indicating a developmental shift in distribution. These findings suggest that AAV9 has preferential tropism for the CNS at earlier gestational stages.

To confirm that E12.5-injected AAV9 provides efficient access to the fetal mouse brain, we repeated the injection comparison (E12.5 vs. E15.5, **Figure 1C**) with AAV9 carrying a CAG-eGFP transgene. Sagittal sections of E15.5-injected embryos revealed biased transgene expression in the heart, liver, lung, with minimal expression in the spinal cord. Closer examination of the brain revealed strong expression in vascular cells, with minimal expression in parenchymal cells. These findings are consistent with previous studies using AAV9 injected at E15 [42]. Embryos injected at E12.5 showed increased brain and spinal cord expression and decreased expression in liver, heart, and lung (**Figure 1H, 1I, Figure S3**). The increased AAV9-mediated CNS expression in E12.5-injected embryos compared to E15.5-injected embryos further supports an age-dependent difference in CNS accessibility, and supports systemic AAV9 delivery at E12.5 as a method for efficient gene delivery to the CNS.

### Midgestational Cre and dual sgRNA delivery enables prenatal CNS genome editing

Cre recombinase is commonly used for conditional knockout of genes, either through viral delivery or Cre-driver transgenic lines. In the context of AAV delivery, Cre can also be used to assess viral tropism at different developmental stages using the Ai14 reporter line [43, 44]. Ai14 reporter mice carry a Cre-dependent tdTomato cassette; delivery of Cre during embryonic development activates tdTomato expression, marking infected progenitor cells and their progeny (**Figure 2A**).

**Figure 2:**
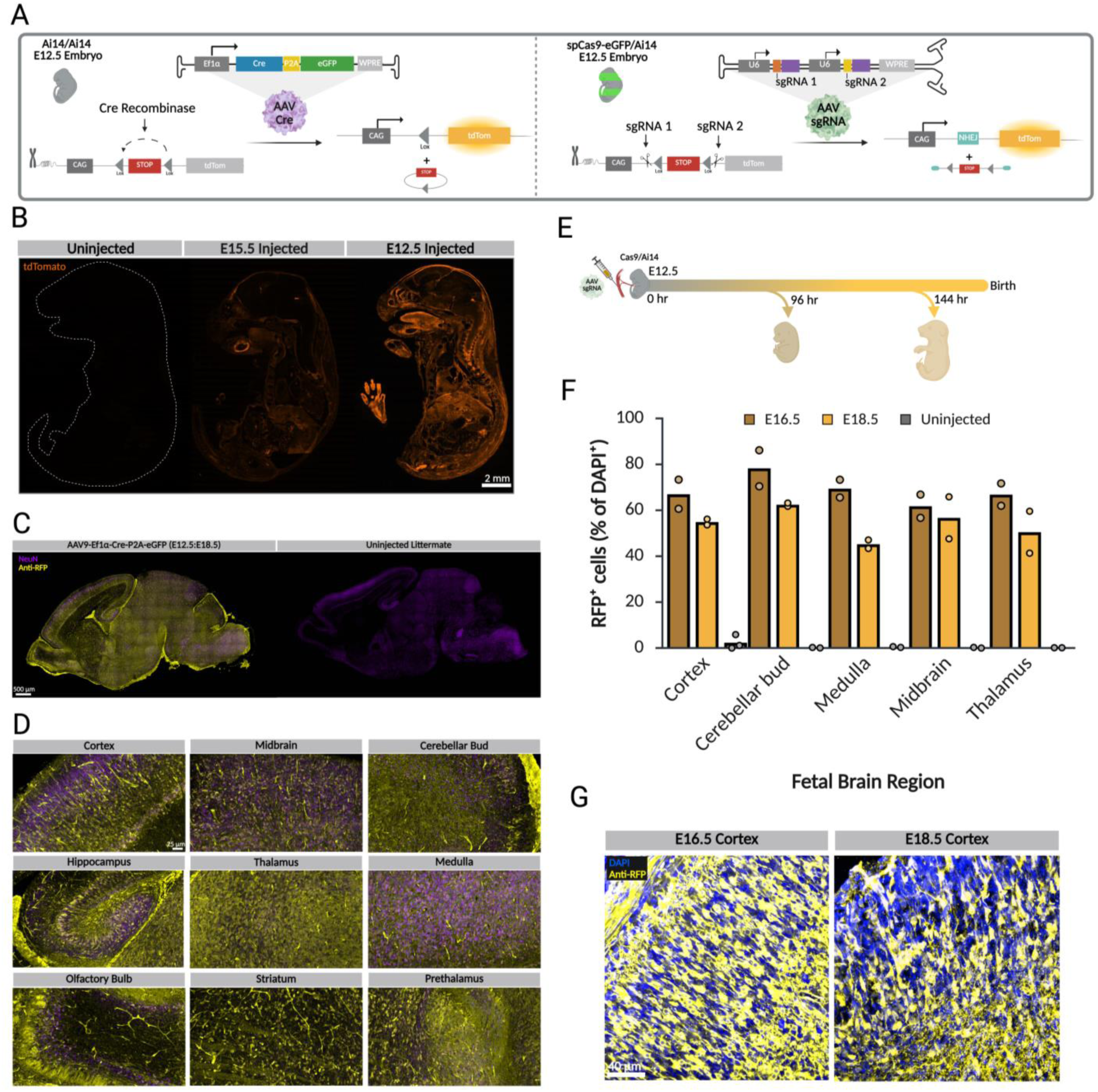
Systemic AAV9-Cre delivery enables prenatal genome recombination in the embryonic brain. **Prenatal genome editing using Cre-Recombinase and CRISPR. A)** Schematic of the Ai14 reporter allele. A loxP-flanked transcriptional stop cassette prevents tdTomato expression from the CAG promoter. In the left panel, delivery of AAV9-Cre excises the stop cassette, enabling tdTomato expression. In the right panel, dual sgRNAs targeting sequences flanking the stop cassette are delivered into Cas9/Ai14 mice, resulting in CRISPR-Cas9-mediated excision of the stop cassette and subsequent tdTomato expression. **B)** Sagittal sections of E18.5 embryos injected with AAV9-Cre at E12.5 or E15.5, showing widespread tdTomato expression across multiple organs. Expression is generally stronger following E12.5 delivery, with noticeable labeling in the brain (n = 3 embryos per stage). **C)** Sagittal sections of P0 brains from injected and uninjected embryos, stained with NeuN and anti-RFP antibodies, showing tdTomato expression in neurons after *in utero* AAV9-Cre delivery (n = 2 embryos per stage). **D)** 25x images of various labeled brain regions from C, illustrating that AAV9-Cre transduced the progenitors of cortical and non-cortical brain areas. **E)** Schematic of in utero injection of dual-sgRNA scAAV9 at embryonic day 12.5 (E12.5; 0 hr). Embryos were harvested at E16.5 (96 hr post-injection) and E18.5 (144 hr post-injection) to assess CRISPR-mediated excision efficiency. **F)** Quantification of RFP-positive cells, expressed as the percentage of DAPI-positive nuclei, across multiple fetal brain regions in uninjected controls and embryos harvested at E16.5 and E18.5 following dual-sgRNA delivery. n = 2 biologically independent embryos per condition. **G)** Representative cortical images showing RFP-expressing cells at E16.5 and E18.5 (40×) following dual-sgRNA injection.

Injection of AAV9-Cre into Ai14 mice at E12.5 and E15.5 led to widespread organ transduction, with the earlier delivery inducing more prominent RFP expression in the CNS (**Figure 2B**). The strong signal in the cortex and other brain regions demonstrates that midgestation viral delivery enables genome manipulation in the embryonic CNS prior to birth (**Figure 2C, 2D**). This supports the potential of this technique for efficient genome editing in the embryonic CNS prior to birth, and suggests that midgestation viral delivery could be used to target neural progenitors for prenatal gene therapy.

To assess the efficacy of CRISPR-mediated genome editing following midgestational delivery, better understand the temporal dynamics of editing, and see how quickly the KO can occur; we evaluated a dual-sgRNA strategy designed to excise the Ai14 floxed stop cassette *in vivo*. Dual-sgRNA in a self-complimentary AAV9 vector was delivered *in utero* at E12.5 into Cas9/Ai14 embryos, and brains were harvested at E16.5 (96 hr post-injection) and E18.5 (144 hr. post-injection) (**Figure 2E**).

Robust RFP expression was observed at both time points, consistent with efficient excision of the stop cassette. Quantification of RFP⁺ cells, expressed as the percentage of DAPI⁺ nuclei across multiple fetal brain regions, revealed substantial reporter activation in injected embryos compared to uninjected controls (n = 2 biologically independent embryos per condition) (**Figure 2F**). Editing was detectable as early as 96 hours post-injection. Representative cortical images demonstrated widespread RFP-positive cells throughout the developing cortex following dual-sgRNA delivery (**Figure 2G**).

### *In utero* conditional knockout of *Reln* via AAV9-Cre induces a prenatal reeler cortex

To further evaluate the potential for *in utero* CNS genome manipulation, we targeted the *Reln* gene for conditional knockout beginning at E12.5. Reelin is a key regulator of cortical lamination, expressed predominantly in marginal zone/layer I (MZ/LI) cells. Loss of *Reln* function results in the reeler cortex, characterized by disrupted laminar organization and, in severe cases, inverted cortical layers [45, 46]. Mice homozygous for a floxed *Reln* exon 1 maintain normal cortical organization, with Cre-mediated recombination inducing a reeler phenotype [47].

Systemic injection of AAV9-Ef1a-Cre-P2A-eGFP into E12.5 homozygous *Reln^flox/flox^* mouse embryos led to displacement of CTIP2-positive layer V neurons and a reduction of Reelin-positive MZ/LI cells at E18.5 (**Figure S4**). This pattern of CTIP2 mislocalization is consistent with prior reports of prenatal Cre-mediated *Reln* knockout and confirms that midgestational AAV9 delivery can be used for genome manipulation in the embryonic CNS. These results support the feasibility of using E12.5-delivered AAV9 to study corticogenesis.

### Midgestational AAV-CRISPR knockout of *Reln* disrupts cortical lamination in wildtype and Cas9-heterozygous mice

Conditional gene deletion using a Cre and floxed sequence relies upon the availability of a suitable mouse line. Programmable nucleases, such as Cas9, have a broad targeting capacity in wildtype animals, thereby increasing the space of sequences that can be assessed. Thus, we assessed whether midgestation delivery of CRISPR-Cas9 machinery to disrupt *Reln* would also yield a reeler-like cortical phenotype. To directly disrupt *Reln* during midgestation, sgRNAs targeting exon 1 of the locus were delivered via AAV9 into E12.5 embryos, which were allowed to develop *in utero* until E18.5. Dual sgRNA delivery into Cas9-expressing embryos, as well as co-injection of a dual sgRNA with SpCas9 into wildtype embryos, resulted in disrupted cortical lamination and a reeler-like phenotype (**Figure 3**). Both approaches caused a strong reduction in RELN-positive cells within the MZ/LI and across the imaged cortical region compared with uninjected controls, consistent with effective loss of functional Reelin protein. Both approaches also showed no reduction in BRN2-positive cells compared to uninjected controls (**Figure 3B**). On average, dual sgRNA delivery and sgRNA/SpCas9 delivery caused a 40- and 9-fold reduction of RELN-positive cells, respectively, compared to control cortices.

**Figure 3:**
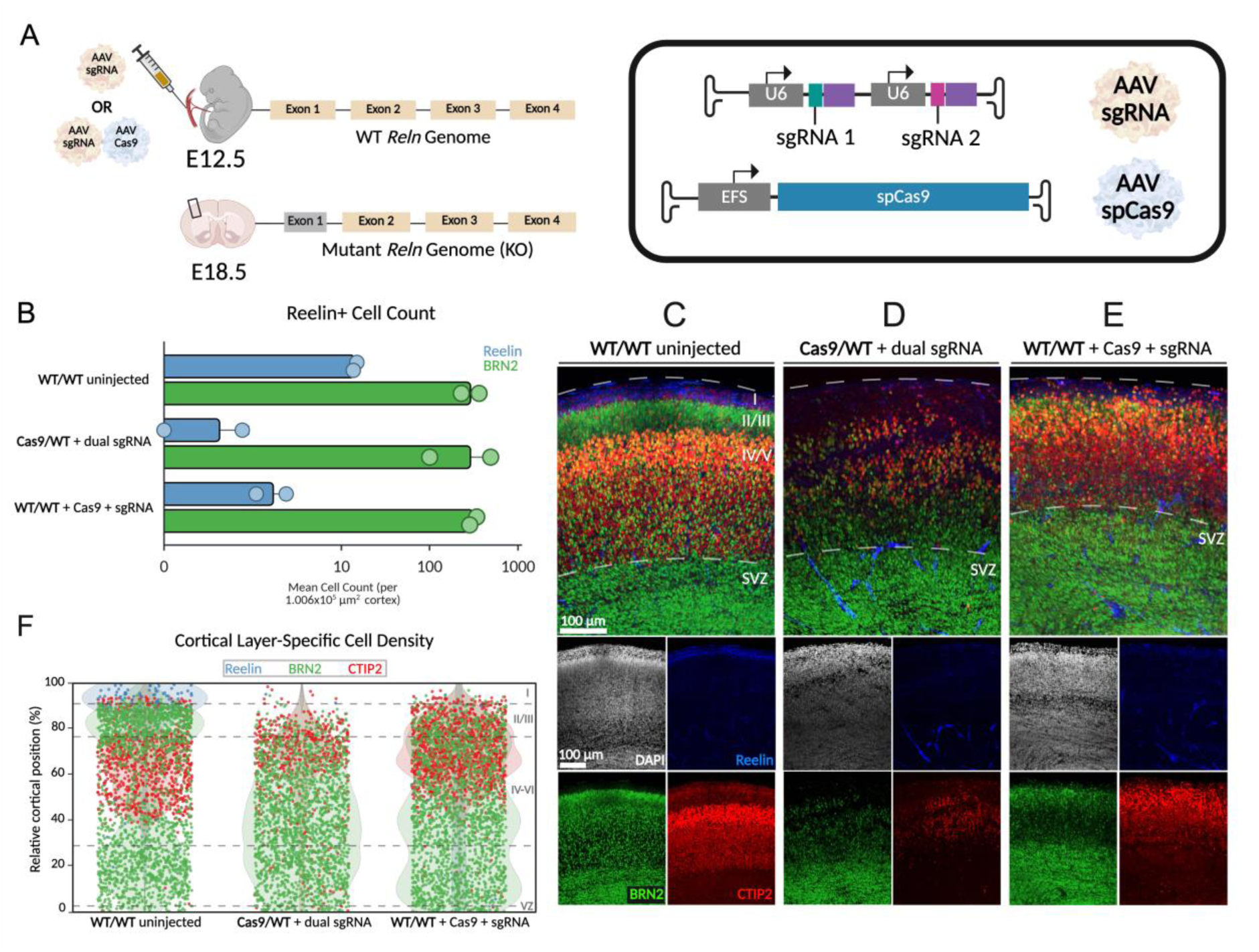
AAV-CRISPR-mediated knockout of *Reln* in midgestation disrupts cortical lamination in wildtype and Cas9-heterozygous mice. **A)** Schematic of AAV-CRISPR microinjection at E12.5. Dual-guide AAV vectors (sgRNA) were delivered into heterozygous Cas9/wildtype (WT) embryos, or co-delivered with an AAV-Cas9 vector (SpCas9 under the EFS promoter) in WT embryos. All transgenes were packaged in an AAV9 capsid. **B)** Quantification of RELN- and BRN2-positive cortical cells at E18.5 following AAV delivery at E12.5 (n = 2 animals per condition). Values are shown on a log scale. Injected conditions exhibit a reduction in RELN-positive cells compared to uninjected wildtype controls. **C-E)** Representative cortical images spanning the subventricular zone (SVZ) to marginal zone (MZ). Layer-specific markers were used for lamination assessment: RELN (LI, blue), BRN2 (LII/III and SVZ, green), and CTIP2 (LIV/V, red). **F)** Relative positions of RELN-, BRN2-, and CTIP2-positive cells plotted as a percentage of total cortical height, illustrating altered cortical organization and reduced RELN-positive cell populations in injected embryos compared to uninjected controls.

Layer-specific markers (Reelin for layer I, BRN2 for layers II/III, and CTIP2 for layer V) were used to evaluate laminar organization. In the absence of Reelin, deeper-layer CTIP2-positive neurons were displaced toward more superficial positions, while normally superficial BRN2-positive neurons were less cohesive and shifted deeper. The Reelin-positive marginal zone was markedly reduced and nearly absent (**Figure 3C-E**). Spatial analysis using strip plots of individual neurons confirmed a reduced density of upper-layer BRN2-positive neurons and mislocalization of deeper-layer CTIP2-positive neurons into superficial regions, highlighting profound laminar disruption following CRISPR-mediated *Reln* knockout (**Figure 3F**). These results demonstrate that midgestational AAV-CRISPR delivery can efficiently manipulate genes in the embryonic CNS, producing functional phenotypes before birth.

### *In utero* HDR-mediated tagging and pathogenic allele installation demonstrate functional precision and broad cortical access

For therapeutic treatment and investigation of the developing embryo, many applications will require precision genome editing. We sought to assess the feasibility of homology-directed repair (HDR) for precise editing in multiple genes of interest. To evaluate access across cortical cell types, we targeted the endogenous *Actb* locus using a previously validated HDR donor and sgRNA [48] (derived from Addgene plasmid #119870) (**Figure 4A**). Midgestational delivery into Cas9 heterozygous embryos induced robust editing across the cortex. HA-positive cells included both SOX2-positive neural stem and progenitor cells and NeuN-positive mature neurons in cortex and hippocampus (**Figure 4B**), indicating that AAV-delivered *in utero* HDR can access a wide range of cortical cell types spanning developmental states.

**Figure 4:**
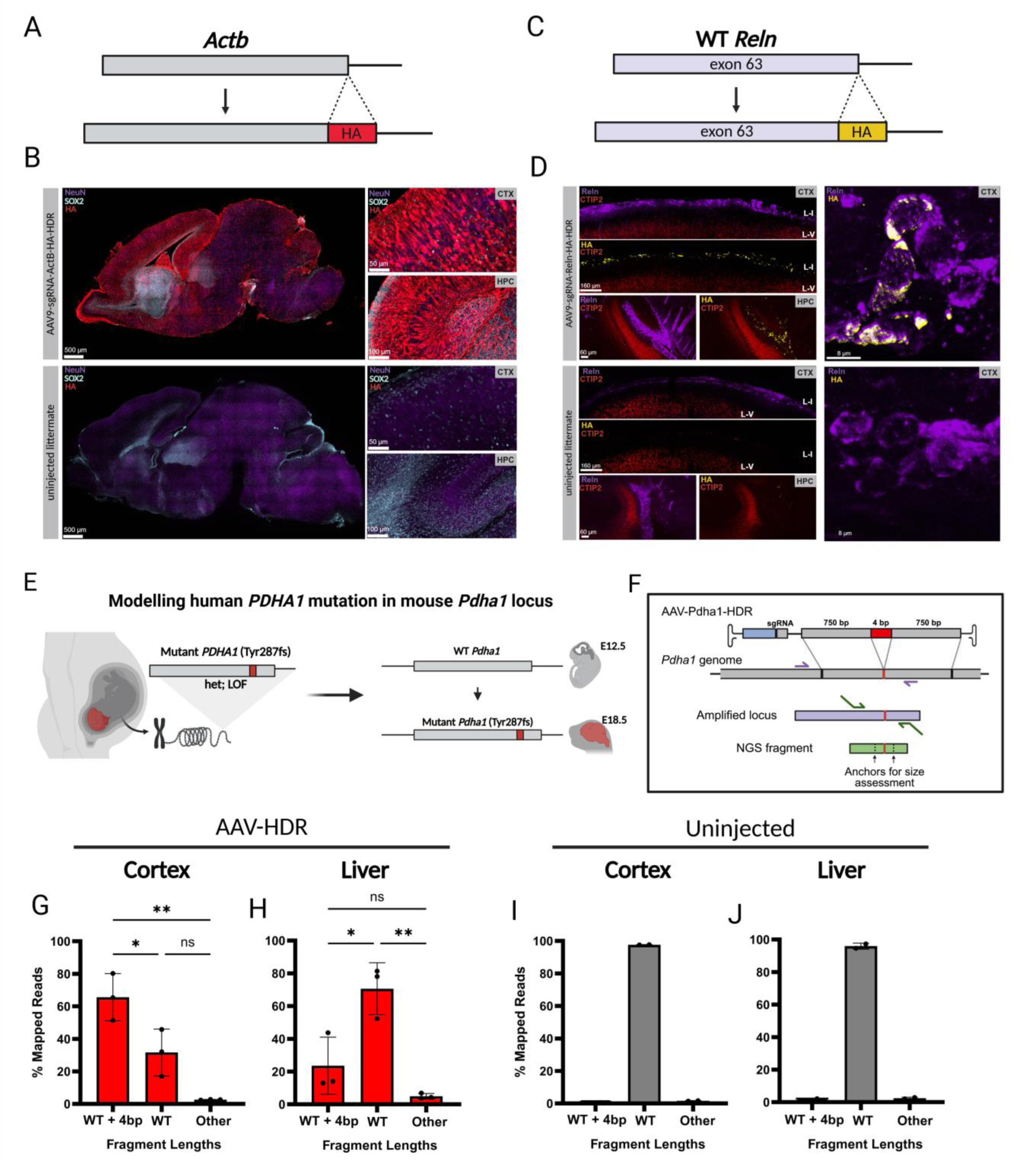
*In utero* HDR-mediated tagging and pathogenic allele installation. **A)** Schematic of HDR-mediated insertion of an HA tag at the C-terminus of WT β-actin (*Actb*). Immunostaining of NeuN (purple), HA (red), and SOX2 (teal) in sagittal brain sections from *Actb*-tagged embryos and uninjected littermates, with higher magnification images of cortex (CTX) and hippocampus (HPC) below. **C)** Schematic of HDR-mediated insertion of an HA tag at the C-terminus of wild-type (WT) *Reln* exon 63, with 750-bp homology arms. **D)** Immunostaining for HA (yellow), RELN (purple), and CTIP2 (red) in E18.5 cortex from Cas9/WT heterozygous embryos. E12.5-injected embryos show HA expression confined to cortical regions (CTX) where RELN is expressed, predominantly in the MZ/LI and hippocampus (HPC). Minimal HA signal was detected in uninjected littermate controls. **E)** Schematic of HDR-mediated incorporation of a human patient-derived PDHA1 Tyr287fs mutation into Cas9/WT heterozygous mouse embryos (WT at the *Pdha1* locus). **F)** AAV transgene HDR design and Pdha1 locus amplification diagram. Sequenced NGS fragments analyzed for HDR incorporation using distance between two anchor sequences within the fragment (64 bp distance between anchors in WT genome). **G-H)** Distribution of reads classified as WT+4bp (HDR 4-bp insertion), WT (no insertion), or “Other” (non-4-bp edits) in injected cortex and liver, plotted as percentage of total reads. HDR insertions significantly exceed WT and other classes (all p < 0.05) in both tissues. (n = 3 embryos). Cortex displays significantly higher insertion frequency than liver (p = 0.0041). **I-J)** Equivalent analysis performed for uninjected cortex and liver (n = 2), showing minimal HDR-like insertion events.

Building on our successful *Reln* cortical knockout, we used HDR to insert an HA epitope at the C-terminus of exon 63 of wild-type *Reln* (**Figure 4C**). This was achieved using a single guide RNA and HA-HDR donor packaged in AAV9. Cortical staining showed colocalization of HA and Reelin, indicating successful genome incorporation (**Figure 4D**). Tagging Reelin did not disrupt cortical lamination, as the locations of layers V and I were preserved relative to uninjected littermates, suggesting that Reelin-HA functions similarly to wild-type Reelin during cortical development. HA-positive cells were detected only in regions where Reelin is endogenously expressed, mainly the marginal zone (LI) and hippocampus.

We next tested whether *in utero* HDR could install a human-derived pathogenic allele. We designed an AAV-HDR transgene encoding the patient-derived PDHA1 Tyr287fs frameshift mutation, which causes a mitochondrial disorder associated with malformations of cortical development (**Figure 4E-F**) [10]. Targeted sequencing confirmed insertion of the 4 bp. HDR frequency was higher in cortex than liver (n = 3, one-way ANOVA, p = 0.0041), and HDR reads were significantly enriched over WT and non-4 bp insertion classes in both tissues (one-way ANOVA with post-hoc correction, all p < 0.05, **Figure 4G-H**). Reads from uninjected littermates were almost exclusively of WT length (n = 2, **Figure 4I-J**).

Taken together, these data show that *in utero* AAV-HDR supports precise and functionally tolerated tagging of endogenous genes, and enables installation of pathogenic human-derived alleles in the developing cortex.

## Discussion

### Establishing midgestational systemic access to the mouse embryo

In this study, we established a platform for systemic *in utero* gene delivery and genome manipulation at midgestational stages in the mouse. By enabling reliable intravenous access to the E12.5 embryo via uterine microdissection, we demonstrated that AAV-mediated gene delivery and precise genome editing can be achieved prior to birth with broad CNS access. This approach defines a previously inaccessible developmental window during which viral delivery, conditional recombination, gene knockout, and HDR-mediated precision editing can be performed in the developing cortex.

A major technical advance of this work is the development of a uterine microdissection strategy that permits direct visualization and access to the vitelline vein at E12.5. Prior systemic *in utero* delivery approaches have largely been restricted to later gestational stages, as the uterus is opaque and densely muscular before E14, limiting visualization of embryonic vasculature [34]. By dissecting a small uterine window directly over the unpaired vitelline vein, we enabled access to the fetal circulation at midgestation. Fluorophore-conjugated dextran injections demonstrated widespread systemic distribution of introduced material to the brain, heart, liver, lungs, and gastrointestinal tract within 30 minutes, confirming effective vascular access at this stage.

### Developmental constraints on AAV capsid performance in the embryo

Using barcoded capsid screening, we found that AAV capsid performance in the embryo is developmentally constrained. At E15.5, engineered AAV9-derived variants previously reported to have enhanced CNS tropism in postnatal or adult settings, including PHP.eB, CAP-B10, and X1.1 [49, 50, 51], did not show significantly enhanced CNS enrichment relative to AAV9 at either the DNA or RNA level. These results indicate that capsid engineering strategies optimized for mature tissues do not necessarily translate to prenatal delivery, likely due to differences in vascular permeability [52], receptor availability, and developmental state.

In contrast, pooled natural serotype screening revealed a pronounced gestational age-dependent shift in AAV9 tropism. When delivered at E12.5, AAV9 demonstrated significantly increased CNS penetrance compared to delivery at E15.5, both at the level of the viral genome and transgene expression. Among eight natural serotypes tested, AAV9 uniquely displayed consistent age-dependent CNS enrichment, accounting for over 70% of normalized viral genomes in the E12.5 brain. Protein-level analysis corroborated these findings, revealing increased brain and spinal cord expression and reduced liver, heart, and lung expression following E12.5 injection. Together, these findings identify midgestation as a permissive developmental window during which AAV9 preferentially accesses the embryonic CNS. Exploiting this window could expand the therapeutic potential of *in utero* gene delivery, particularly given that existing approaches are frequently constrained by broad peripheral transduction. Enhancing CNS specificity while minimizing off-target organ expression represents a critical objective for advancing prenatal gene therapy strategies. Combining capsid selection and midgestational systemic delivery with cell-type specific promoters, miRNA target sites, and other regulatory elements would likely increase specificity and could largely limit expression to the CNS.

### Efficient prenatal genome manipulation using Cre and CRISPR-based approaches

Leveraging this permissive window, we demonstrated efficient genome manipulation in the embryonic CNS using both Cre-lox and CRISPR-based strategies. Systemic AAV9-mediated delivery of Cre recombinase to Ai14 reporter embryos resulted in widespread recombination across multiple organs by E18.5, with particularly high penetrance in the brain following E12.5 administration. Because recombination affects both infected progenitors and their progeny, this approach provides a rapid and scalable alternative to Cre driver lines for CNS-specific genome manipulation prior to birth [53].

Importantly, the dual-sgRNA strategy enables deletion of exon-sized genomic fragments through Cas9-mediated excision between two guide RNA target sites. This approach extends prenatal genome manipulation beyond Cre-lox recombination by permitting precise programmable gene disruption in the embryonic CNS. The ability to induce defined deletions in neural progenitors during midgestation supports the feasibility of in utero CRISPR-based gene knockout strategies and provides a framework for modeling neurodevelopmental disorders or exploring prenatal therapeutic genome editing.

Beyond reporter systems, we demonstrated functional genome disruption by targeting *Reln*, a gene essential for cortical lamination [54]. Both conditional deletion of *Reln* via AAV9-Cre delivery as well as AAV-CRISPR-mediated knockout initiated at E12.5 were sufficient to induce a prenatal reeler-like phenotype. Edited embryos exhibited loss of Reelin-positive marginal zone cells, redistribution of deep-layer neurons, and disrupted cortical lamination consistent with classical reeler pathology [55]. These findings establish that midgestational genome manipulation can perturb core neurodevelopmental programs prior to birth in otherwise wild-type animals.

A practical advantage of this platform is the use of Cas9-expressing stud males crossed with wild-type females, producing embryos with heterozygous Cas9 expression. As the pregnant females were terminal, this strategy is both cost-effective and scalable, facilitating efficient generation of somatically edited embryos without the requirement for maintaining complex transgenic breeding colonies. While germline editing is possible, it was not systematically assessed in this study and remains an important area for future evaluation.

### Precision genome modification via HDR in the embryonic CNS

In addition to gene disruption, we demonstrated the feasibility of precise genome modification via HDR *in vivo*. HA tagging of endogenous *Reln* preserved protein localization and cortical lamination, indicating that C-terminal tagging did not compromise Reelin function during corticogenesis. Extension of this strategy to the *Actb* locus revealed robust and widespread cortical access, with successful HA knock-in observed in neural progenitors and differentiated neurons across the cortex and hippocampus. Precise editing by HDR is particularly well suited to this developmental context, as the E12.5 cortex is dominated by rapidly dividing neural progenitors that have more capacity for HDR [56, 57]. Together, these results establish HDR-based editing as a viable strategy for endogenous gene tagging and disease modeling in the embryonic CNS.

### Modeling prenatal cortical disorders and clinical relevance

The ability to manipulate the embryonic genome systemically and assess phenotypes prior to birth has direct relevance for modeling congenital neurodevelopmental disorders. Many pathogenic variants identified through prenatal screening, particularly those associated with malformed cortical development, exhibit variable penetrance, and uncertain postnatal manifestation [58]. The majority of prenatally diagnosed cortical malformations are associated with single nucleotide or copy number variants [10], underscoring the need for precise functional modeling. Guided by these observations, we modeled a human pathogenic four-base-pair insertion in the PDHA1 locus associated with prenatal malformation of cortical development, using HDR-mediated editing. This approach enables recapitulation of disease-associated alleles *in vivo* and allows direct assessment of developmental consequences during corticogenesis. *In utero* modeling could provide a powerful framework for phenocopying early human neurodevelopment prior to term, offering insight into how specific genetic lesions may translate into postnatal disability.

### Gestational equivalence and translational interpretation

Direct comparison between mouse and human gestational timelines remains challenging due to species-specific differences in brain development. Rather than relying on global embryonic age, we define gestational equivalences using cortex-specific developmental and neurogenesis milestones. Using this framework, mouse E12 corresponds approximately to human gestational weeks (GW) 7-10, characterized by early neurogenesis and the birth of deep-layer neurons [59, 60]. Mouse E15 aligns with GW10-14, a period of peak deep-layer neurogenesis and expanding progenitor populations, while mouse P0 approximates GW20-26, when cortical layers are established and gliogenesis begins [61, 62]. Thus, the midgestational mouse interventions we describe here offer translational relevance for modeling early human cortical development.

### Technical challenges and sources of experimental variability

Despite the advances we describe here, *in utero* gene delivery remains technically challenging. We observed embryonic lethality associated with excessive uterine manipulation, prolonged exposure leading to tissue desiccation, uterine contortion, or obstruction of placental blood flow. Creation of an injection site that was too large resulted in embryonic inviability, likely due to loss of amniotic integrity or hemorrhage. Maternal morbidity also posed a significant limitation. Dams that failed to carry embryos to term often exhibited near-necrotic intestines and marked fecal accumulation at necropsy, suggestive of ileus or intestinal damage incurred during surgery. These findings emphasize the importance of minimizing operative duration and tissue disruption.

A further source of variability in our experiments arose from embryonic staging. Although injections were performed at midday on E12 based on plug detection, we observed apparent developmental stages ranging from E11.5 to E13.5, likely reflecting variation in copulation timing during the nocturnal active phase and differences in embryo number. Given the stage-dependent nature of cortical development and genome editing efficiency, more precise methods of tracking gestational time would aid future experiments.

### Summary and Future directions

In summary, this work establishes a versatile framework for systemic *in utero* gene delivery and genome editing at midgestation in the mouse, enabling gene disruption, conditional recombination, and precision editing within a permissive developmental window. By extending access to the embryonic circulation at E12.5, we provide a powerful platform for modeling congenital neurological disorders and exploring early therapeutic interventions prior to birth.

## Materials and Methods

### Plasmid DNA

Standard molecular cloning techniques were used for plasmid DNA production and assembly. Double-stranded DNA fragments were synthesized by Twist Bioscience or Integrated DNA Technologies and inserted into pAAV backbones using NEBuilder HiFi DNA Assembly (New England Biolabs, E2621). sgRNA sequences were synthesized as overlapping single-stranded DNA fragments, annealed, and ligated into expression cassettes using T4 DNA Ligase (New England Biolabs, M0202). Self-complementary pAAVs were generated from pscAAV-CAG-GFP, a gift from Mark Kay (Addgene, 83279).

pAAV2/1 (gift from James M. Wilson; Addgene plasmid #112862), pAAV2/2 (gift from Melina Fan; Addgene plasmid #104963), AAV3 rep-cap (Cell Biolabs, VPK-423), AAV4 rep-cap (Cell Biolabs, VPK-424), pAAV2/5 (gift from Melina Fan; Addgene plasmid #104964), AAV6 rep-cap (Cell Biolabs, VPK-426), pAAV2/7 (gift from James M. Wilson; Addgene plasmid #112863), pAAV2/8 (gift from James M. Wilson; Addgene plasmid #112864), pUCmini-iCAP-AAV-PHP.eB13 (Addgene #103005), pUCmini-iCAP-AAV.CAP-B10 (Addgene #175004), AAV-DJ rep-cap (Cell Biolabs, VPK-420-DJ), and pUCmini-iCAP-AAV9-X1.1 (Addgene #196836) were used for production of AAVs together with the adenoviral helper plasmid pHelper (Agilent Technologies, #240071).

### AAV Production

AAV vectors were produced following established protocols available on protocols.io (https://doi.org/10.17504/protocols.io.n2bvjnew5gk5/v1; https://doi.org/10.17504/protocols.io.e6nvw1n47lmk/v1). HEK293T cells (ATCC, CRL-3216; RRID: CVCL_0063) were triple-transfected with the Rep-Cap plasmid, the genome packaging plasmid, and the pHelper using PEI-MAX (Polysciences, 24765). Viral particles were harvested from cell lysate and supernatant and purified using iodixanol gradient ultracentrifugation (15%, 25%, 40%, 60%) in a Type 70 Ti rotor (Beckman Coulter, 337922) at 58,400 rpm for 1.5 hours. Viral preparations were buffer-exchanged and concentrated using Amicon Ultra-15 centrifugal filters (100 kDa MWCO; MilliporeSigma, UFC9100) into sterile DPBS (Thermo Fisher Scientific, 14190144). Viral titers were quantified by qPCR using DNase I-resistant viral genomes relative to a linearized plasmid standard. AAVs were diluted in sterile saline before *in vivo* use. All non-pooled injections were at a dose of 2×10^11^ vg, unless specified otherwise.

### Cell Culture

HEK293T (ATCC, CRL-3216; RRID: CVCL_0063) and NIH3T3 (ATCC, CRL-1658; RRID:CVCL_0594) cells were maintained in DMEM (Thermo Fisher Scientific, 10569010) supplemented with 10% FBS (Cytiva, SH30070.03). NIH3T3 cells were seeded at 75% confluence for HDR assay validation. Transfections were performed using Lipofectamine LTX (Thermo Fisher Scientific, 15338100) with DNA amounts adjusted to minimize toxicity.

### Animal Husbandry

All procedures were approved by the Institutional Animal Care and Use Committee and Office of Laboratory Animal Resources at the California Institute of Technology (protocol #1746) and followed the NIH Guide for the Care and Use of Laboratory Animals. Breeding-age male and female C57BL/6J (JAX #000664, RRID: IMSR_JAX:000664), Ai14 (JAX #007914, RRID: IMSR_JAX:007914), Rosa26-Cas9-eGFP (JAX #026179, RRID: IMSR_JAX:026179), and Reln-Flox (JAX #037348, RRID: IMSR_JAX:037348) mice were housed under a 12-h light/dark cycle with *ad libitum* access to food and water. Females were group-housed (3-5 per cage) and males singly housed as studs.

### Timed Matings

Female mice were placed into stud male cages during the active phase. A copulatory plug was taken as evidence of mating, and 12 PM of the plug-detection day was designated as embryonic day 0.5 (E0.5).

### Surgery and Injections

Mice were anesthetized with isoflurane (5% for induction and 1% for maintenance). Animals were then given subcutaneous ketoprofen (5 mg kg⁻¹), buprenorphine XR (3.25 mg kg⁻¹), and 1 mL sterile saline. The surgical area was sterilized, and 3 drops of 0.5% bupivacaine were injected subcutaneously at the incision site for local anesthesia.

#### Laparotomy and Uterine Window

At E12.5, a 2.5 cm laparotomy was performed to expose the uterine horns. Individual horns were gently exteriorized with sterile forceps, wrapped in saline-soaked sterile gauze, and manipulated to visualize single embryos. Opposite the placental implantation site, and above clustered blood islands, the uterine wall was dissected with fine tweezers to create a ∼3.5 mm window exposing the vitelline vein on the yolk sac surface (Supplementary Video 1, https://doi.org/10.6084/m9.figshare.31516642).

#### Needles and Microinjection

Glass capillary tubes (G100TF-4, OD 1.00 mm; Warner Instruments) were pulled on a vertical needle puller, cut using a microforge (MF2; Narishige) to a ∼20 µm tip diameter, and beveled at 20° using a microgrinder (EG-402; Narishige). Concentrated AAV or dextran dye (Thermo Fisher Scientific, D1817) was loaded into needles and mounted on a microinjector (FemtoJet 4i, Model 5252000021; Eppendorf). Under a stereoscope, the needle was positioned parallel to the vitelline vein and advanced until fetal red blood cells flashed, indicating successful entry. Approximately 5 µL of viral solution was pulse-injected into the direction of blood flow. After microinjection of 3-5 embryos, the uterine horns were returned to the abdominal cavity, the incision was sutured closed, and dams were maintained for 3-6 days.

### Embryo Collection and Tissue Processing

At E18.5, dams were euthanized and injected embryos identified by relative uterine position. Embryos were decapitated and placed in warm saline on a shaker for 15-20 min for passive perfusion. For peripheral organs (liver, lungs, heart, gonads), tissues were dissected and fixed in 4% paraformaldehyde (PFA) in PBS for 24 hours. For brain fixation, skin and eyes were removed, and brains were fixed in-skull in 4% PFA for 48 hours.

### Pooled Injections

AAV pools were constructed with unique 8-nt barcodes placed in a noncoding region of each transgene. Viral preparations were pooled in sterile saline to final concentrations of 1.6×10¹⁰ vg/variant/5 µL (natural serotype comparison) and 4.4×10⁸ vg/variant/5 µL (engineered capsid pool). At time of injection, aliquots of each virus pool were frozen for subsequent sequencing normalization.

### NGS and Analysis

Viral and genomic DNA were isolated and prepared for sequencing using a modified protocol for pooled capsid screens (https://doi.org/10.17504/protocols.io.bp2l695zklqe/v2). Briefly, ∼50 mg tissue was homogenized in TRIzol (Life Technologies, 15596) using a BeadBug homogenizer (Benchmark Scientific, D1036). Following the episomal DNA extraction, the transgene was PCR-amplified in three rounds (∼500 bp primary; ∼180 bp variable region; NEBNext Dual Index primers; New England Biolabs, E7600). Products were gel-purified (Zymo Zymoclean Gel DNA Recovery Kit, D4021) and quantified via Qubit. Indexed libraries were sequenced on an Illumina HiSeq 2500 (Millard and Muriel Jacobs Genetics and Genomics Laboratory, Caltech).

Sequencing data were processed in Python 3.11 (Google Colab). Reads were mapped to a reference barcode library using Biopython (v1.86) and numpy, and barcode counts were tabulated using pandas. For E15.5 pooled screens, counts were normalized to barcode frequencies in the input viral pool to compute tissue-specific enrichment. Data visualizations were generated using matplotlib. Genomic DNA for HDR validation was extracted using the DNAeasy Blood & Tissue Kit (Qiagen, Cat. No. 69504). Length analysis was performed from mapped FASTQ data using a custom Python pipeline that located start/end anchors (allowing up to one mismatch) and plotted fragment length distributions using GraphPad Prism.

### Sectioning and Immunohistochemistry

Tissue was sectioned using either a vibratome (65 µm thick sections) or a cryostat (whole embryos, sagittal sections, 50 µm thick). Sections were permeabilized for 1 hour at room temperature in PBS containing 0.5% Triton X-100, followed by blocking for 1.5 hours at room temperature or overnight at 4 °C in PBS with 0.1% Triton X-100 and 10% normal donkey serum. Primary antibodies were applied overnight at 4 °C in blocking solution at 1:500-1:1000 dilution and included: Anti-Reelin (Sigma, MAB5364), Anti-Ctip2 [25B6] (Abcam, ab18465), Chicken Anti-GFP (Aves Labs, GFP-1020), Anti-RFP (Rockland, 600-401-379), Brn2/POU3F2 (Cell Signaling, mAb 12137), Anti-Sox2 (Abcam, ab79351), and Anti-NeuN (Abcam, ab279297). Sections were washed three times for 10 minutes in PBS and incubated with species-appropriate secondary antibodies at 1:200 dilution for 1.5 hours at 4 °C: Goat anti-Chicken IgY Alexa Fluor 488 (Thermo Fisher, A32931), Donkey anti-Rat IgG Alexa Fluor 594 (Thermo Fisher, A21209), Donkey anti-Rat IgG Alexa Fluor 647 (Abcam, ab150155), Donkey anti-Mouse IgG Alexa Fluor 555 (Thermo Fisher, A32773). After secondary incubation, sections were washed for 30 minutes with DAPI (1:5,000 in PBS; Thermo Fisher, 62248) at room temperature, followed by two additional 10-minute washes in PBS. Stained sections were mounted using ProLong Diamond Antifade Mountant (Thermo Fisher, P36970) and imaged using a spinning disk confocal microscope as described in the Imaging and Analysis section.

### Imaging and Analysis

Images were acquired using a spinning disk confocal microscope (Dragonfly; Andor) with Fusion software and 25× water- or 40× oil-immersion objectives (Leica). Zyla sCMOS cameras (Andor) were used for image capture. Three coronal sections spanning from the subventricular zone (SVZ) to the marginal zone (MZ) were sampled per animal for Reelin KO quantification. Five z-planes were counted per section, and means were plotted per animal. Image preprocessing was performed in ImageJ/Fiji. Automated segmentation and counting were performed using a threshold-based Python pipeline (scikit-image, scipy, numpy, pandas). TdTomato-positive cells were quantified in Cas9/Ai14 heterozygous embryos using a Python threshold-based pipeline. Z-stacks (16 planes, 2 µm spacing) were max intensity projected, DAPI nuclei segmented and expanded 5 pixels, and background-corrected TdTomato intensity per cell was measured; cells above 75 a.u. were counted, and percent positivity calculated. Anatomical regions were assigned using the Allen Institute for Brain Science Developing Mouse Brain Atlas (https://developingmouse.brain-map.org) [36].

### Statistics and Reproducibility

Sample sizes are reported in figure legends. No samples or data were excluded. Statistical tests are indicated in figure legends. Analyses were performed in GraphPad Prism (version 10.6.1). NS, not significant (p>0.05); *p < 0.05, **p < 0.01, ***p < 0.001 ****p< 0.0001.

## Acknowledgments

We thank the members of the Gradinaru laboratory for thoughtful feedback and helpful discussions throughout the development of this work, with special thanks to C. Oikonomou for careful review of the manuscript. We also thank P. Anguiano for outstanding administrative support. This work was supported in part by the Gates Foundation (Award No. INV-077908 to V.G). The conclusions and opinions expressed in this work are those of the authors alone and shall not be attributed to the Foundation. Under the grant conditions of the Foundation, a Creative Commons Attribution 4.0 License has already been assigned to the Author Accepted Manuscript version that might arise from this submission. Please note works submitted as a preprint have not undergone a peer review process. This work was also supported in part by the NIH Pioneer Grant (Award No. 5DP1NS111369 to VG). Schematics were created using BioRender.

## Author Contributions

C.R.J. conceived the study, oversaw the project direction, wrote the manuscript, and generated the figures with input from all authors. V.G., G.C., and M.B. assisted with manuscript editing and preparation. C.R.J. carried out the majority of experiments and performed data collection and analysis. C.R.J and M.B. conceptualized and optimized microinjection techniques. Carrie R.J. contributed to histology experiments and reeler cortical phenotype assessment. N.A. supported molecular cloning and viral production. M.B. and G.C. assisted in CRISPR/HDR construct designs. V.G. supervised the project and provided funding.

---

**Figure S1:**
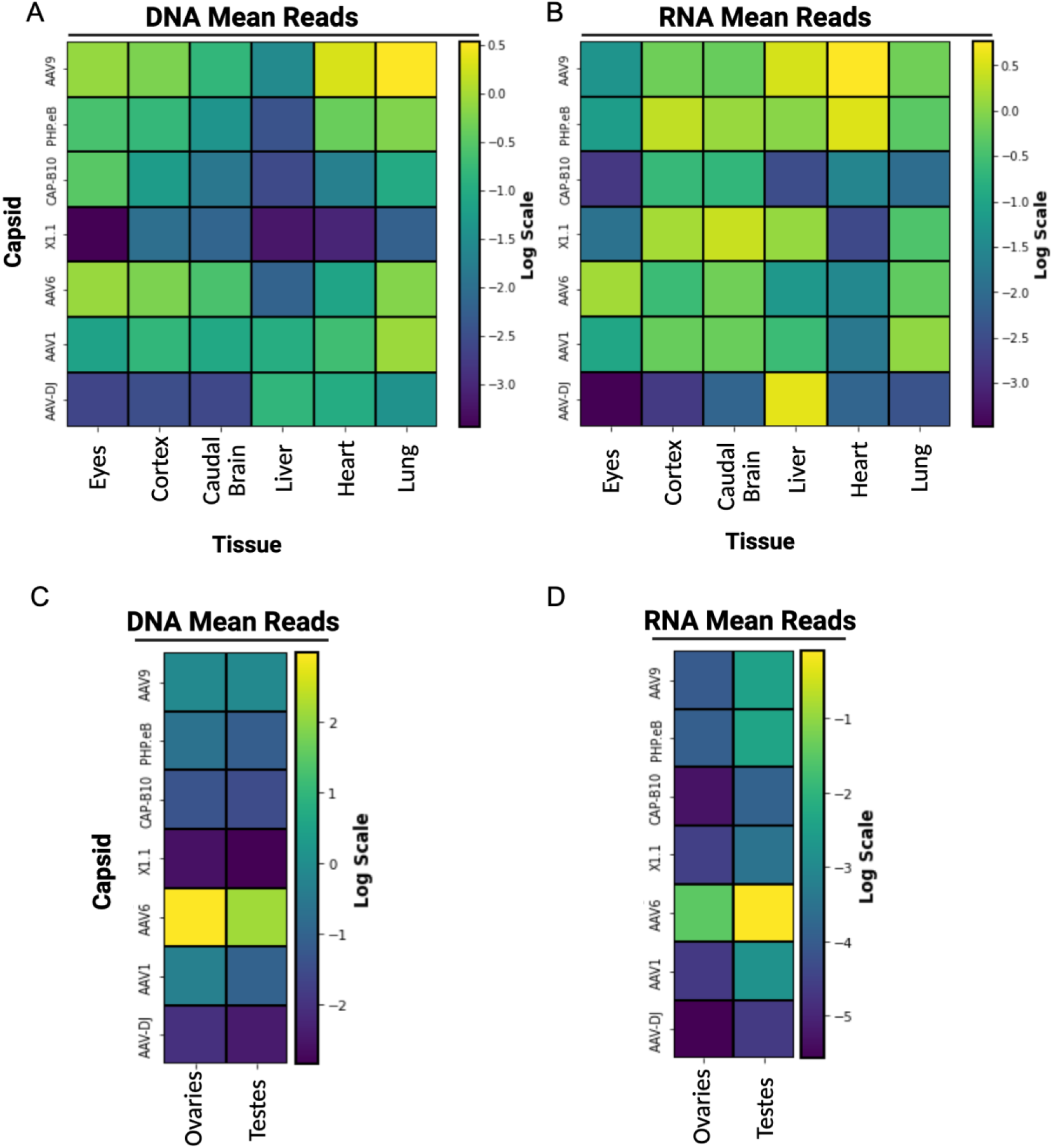
Barcoded AAV capsid screening at E15.5 reveals no enhanced CNS tropism over AAV9. Eight embryos were injected *in utero* at E15.5 with a pooled barcoded library of seven capsids (AAV9, AAV-DJ, AAV1, AAV6, CAP-B10, PHP.eB, X1.1) at equal concentrations and collected at E18.5. Data were analyzed using linear mixed effects models with tissue, capsid, and normalized reads as fixed effects, and biological replicates as a random effect, with AAV9 set as the reference capsid. **A-B)** Normalized DNA **(A)** and RNA **(B)** abundance of each capsid in cortex, caudal brain, eyes, liver, heart, and lung, relative to AAV9, quantified by NGS. Brain-biased capsids (PHP.eB, CAP-B10, X1.1; tropism established in adult mice) showed no statistically significant enrichment in CNS tissues or depletion in liver (all p > 0.05). **C-D)** Normalized DNA **(C)** and RNA **(D)** abundance in gonads (ovaries and testes; n = 2 female, n = 6 male).

**Figure S2:**
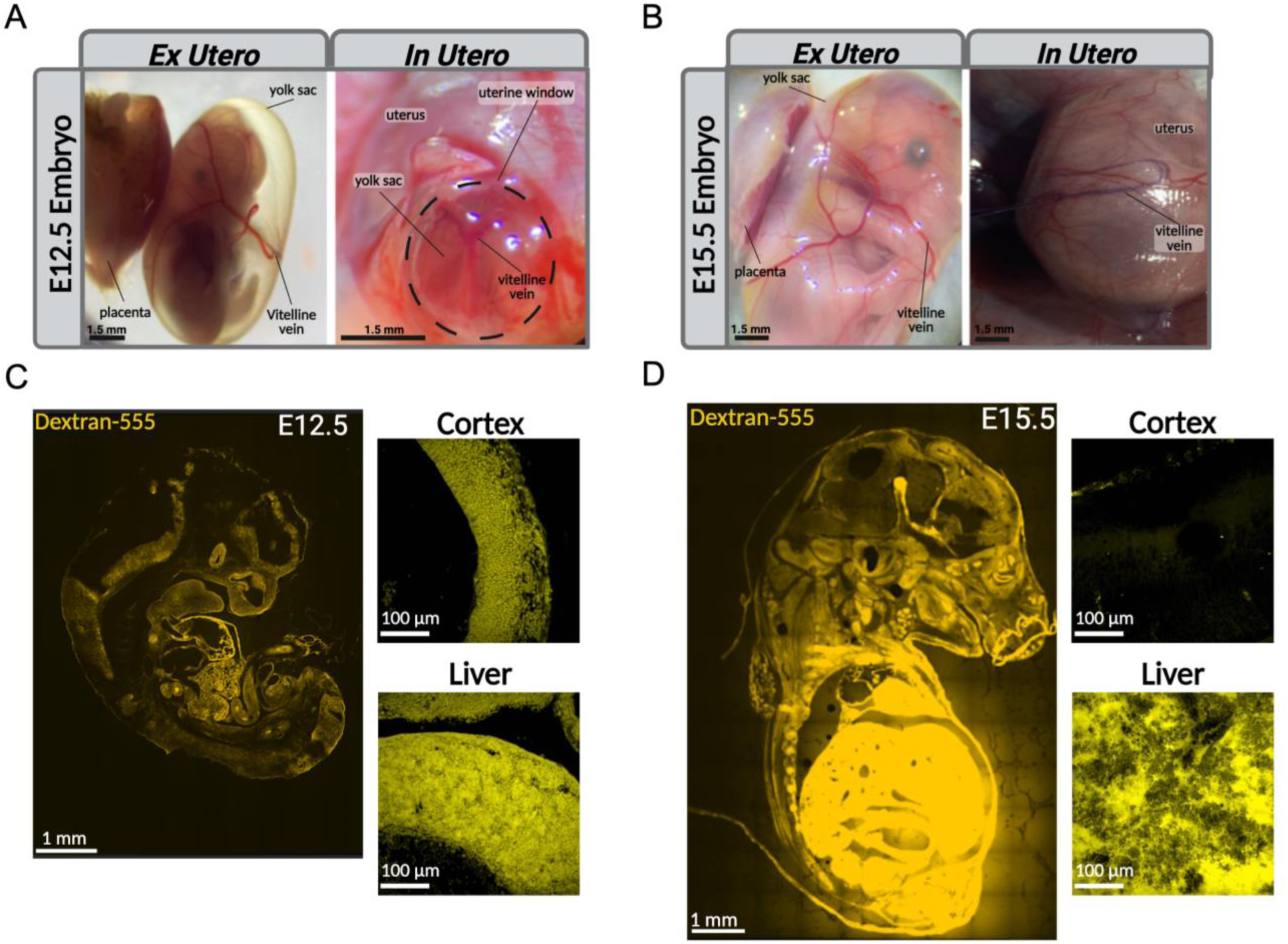
Systemic access via vitelline vein injections at E12.5 and E15.5. **A-B)** *In utero* and *ex utero* views of E12.5 (A) and E15.5 (B) embryos, showing yolk sac and uterine anatomy. E12.5 anatomical images are repeated from Figure 1C. **C-D)** Fluorophore-conjugated dextran dye (555 nm, yellow) was injected through the vitelline vein. Embryos were collected within 1 hour of injection. Broad systemic distribution was observed in brain, heart, liver, lungs, and gut. Magnified images of E15.5 cortex and liver highlight dye uptake in organs of interest. Fluorescence intensity was normalized to liver intensity within each embryo (n = 2 embryos per stage)

**Figure S3:**
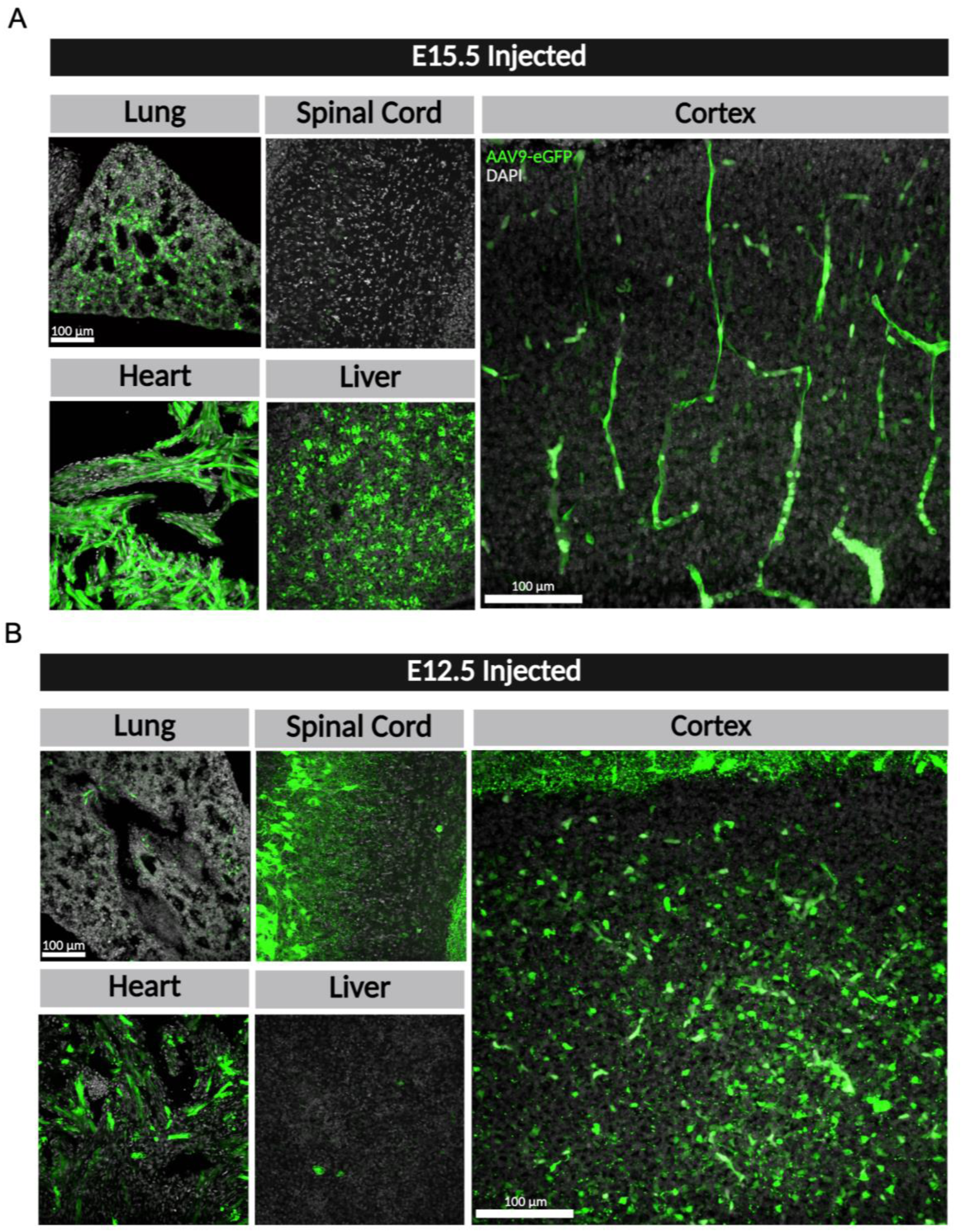
Age-dependent AAV9 transgene tropism in E15.5- and E12.5-injected embryos. **A)** Representative images of E18.5 embryos injected at E15.5 with AAV9-eGFP showing transgene expression in heart, liver, and lung with minimal signal in cortex and spinal cord. **B)** Representative images of E18.5 embryos injected at E12.5 with the same AAV9-eGFP construct showing increased expression in cortex and spinal cord and reduced expression in heart, liver, and lung. Images highlight the developmental shift in AAV9 tropism between E12.5 and E15.5 embryos.

**Figure S4:**
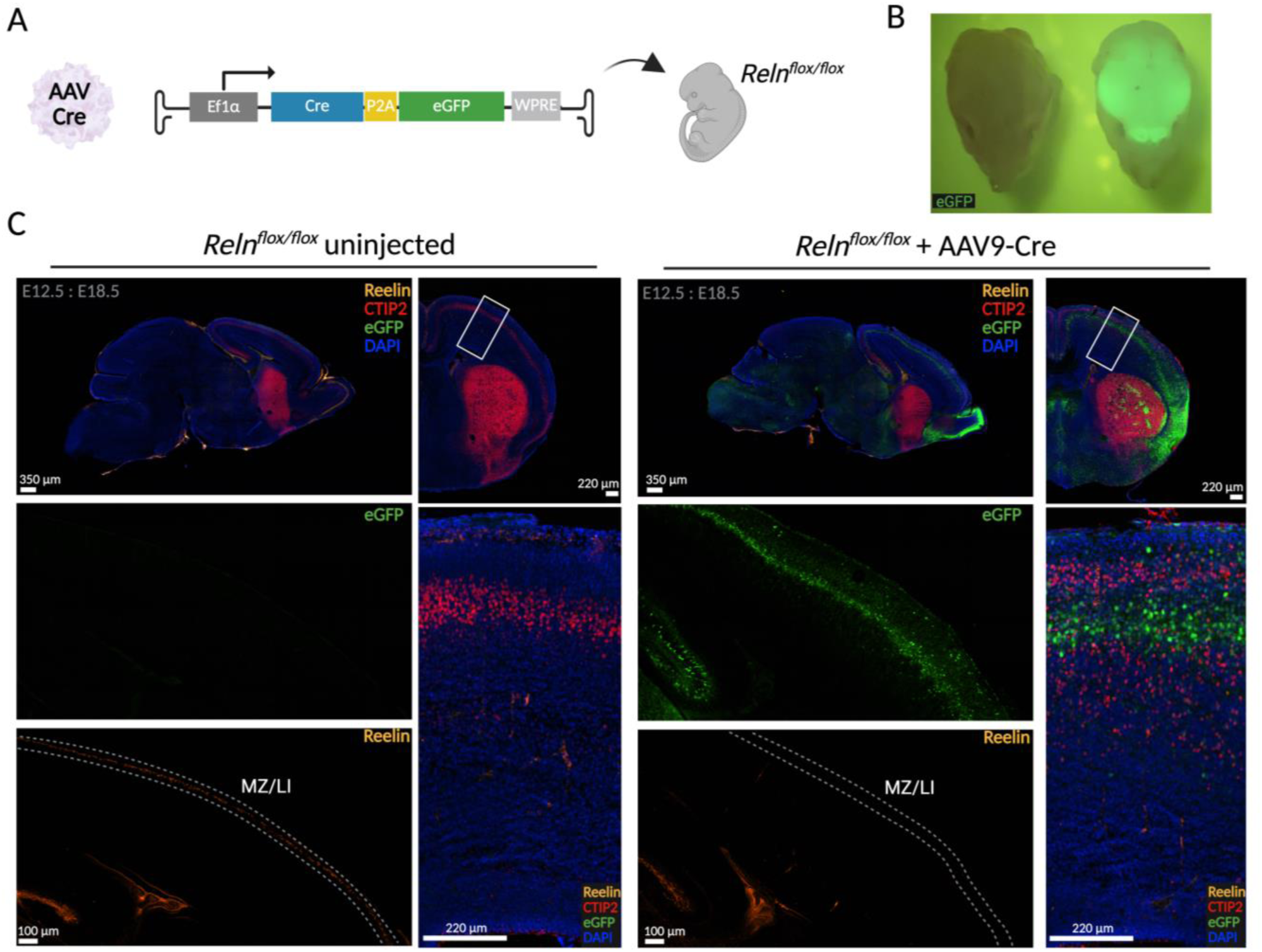
AAV-Cre-mediated prenatal induction of reeler cortex. **A)** Schematic of injection of AAV9 carrying a Cre transgene into E12.5 homozygous *Reln^flox/flox^*embryos. **B)** Brain transduction by Cre-P2A-eGFP transgene at E18.5. Uninjected (left) and injected (right) brains were imaged *in situ* within the skull. **C)** Sagittal E18.5 brain sections showing eGFP and immunostaining for RELN and CTIP2. AAV-Cre injection leads to displacement of deep-layer CTIP2-positive neurons and loss of RELN-positive marginal zone/layer I cells, consistent with prenatal reeler cortex.

